# Clinical Metabolomics studies in diagnosing Prediabetes and Type 2 Diabetes: Unveiling Trends through Bibliometric Analysis

**DOI:** 10.64898/2026.02.28.708687

**Authors:** Mengmeng Li, Mengmeng Sun, Mengyuan Li, Lin Yao, Wenzhe Wang, Jian Mei, Xupeng Huang, Xinyu Zhang, Zheng Lian, Thanh Tuan Vu Le, Min He, Hongfeng Wang

## Abstract

**Background:** Prediabetes is the transitional stage preceding the onset of type 2 diabetes. Due to its asymptomatic nature and slow progression, conventional screening methods frequently fail to identify individuals in this prediabetic stage. Therefore, this study employs bibliometric methods to reveal the abnormal changes in clinically relevant metabolites in diabetes, providing evidence for early diagnosis, pathogenesis, and precision interventions.

**Methods:** Search the Web of Science Core Collection database for all papers on metabolomics related to prediabetes and type 2 diabetes clinical research from the database’s inception until May 2025. Use VOSviewer and CiteSpace software to perform bibliometric analysis on the included literature.

**Results:** A total of 1,742 studies were ultimately included in the review of the literature. Researchers from China, the United States, and Germany, as well as institutions, have made significant contributions to metabolomics research in prediabetes and type 2 diabetes. The highest number of publications and citations was from the journals Metabolites and Diabetes. The current research hotspots in this field, as determined by keyword co-occurrence and clustering analysis, include metabolomics, insulin resistance, obesity, risk factors, biomarkers, gut microbiota, amino acids, lipidomics, and mass spectrometry.

**Conclusion:** The outcomes of the study are instrumental in enabling scholars to comprehend the developmental trajectory of metabolites associated with prediabetes and type 2 diabetes and facilitate the rapid identification of emerging research pathways.

## 1 Introduction

Type 2 diabetes (T2D) is a complex, progressive metabolic disorder characterized by abnormal hepatic gluconeogenesis^1^, pancreatic β-cell dysfunction, and peripheral insulin resistance (IR) leading to impaired glucose metabolism^2,3^. The core diagnosis of this disease is primarily based on the presence of hyperglycemia, which is associated with an elevated risk of both microvascular and macrovascular complications^4^. According to the International Diabetes Federation (IDF), the global diabetes population surpassed 460 million in 2019 and is projected to reach 700 million by 2045^5^. It is worth noting that T2D has a long asymptomatic stage, during which blood glucose levels are already impaired but have not yet reached the diagnostic criteria for T2D^4^. This state is termed prediabetes (Pre-DM)^6^. Pre-DM not only represents a critical risk stage for T2D development, but it also dramatically raises the risk of cardiovascular and cerebrovascular disease^6^. According to current data, the prevalence of Pre-DM among adults in the United States is 38%, whereas in China it is 35.7%^7^. This indicates that T2D and Pre-DM have become one of the major challenges in the field of global public health.

According to the diagnostic criteria established by the World Health Organization (WHO, 1999), Pre-DM is defined based on the oral glucose tolerance test (OGTT) and encompasses impaired fasting glucose (IFG), impaired glucose tolerance (IGT), as well as a combination of both states (IFG + IGT)^7^. Although all three conditions fall under the category of Pre-DM, IFG primarily manifests as hepatic IR and insufficient basal insulin secretion, whereas IGT is characterized by postprandial hyperglycemia resulting from muscle IR^8,9^. Additionally, clinical research has revealed that only a minority of individuals with Pre-DM fulfill the diagnostic criteria for all three of the subtypes above concurrently^10^. Although the OGTT is considered the gold standard for diagnosing Pre-DM and T2D, its requirement for multiple blood samples over a two-hour period and its complex, time-consuming procedure limit its widespread adoption in routine clinical practice^11^. Therefore, current clinical practice often employs fasting plasma glucose (FPG) and/or glycated hemoglobin (HbA1c) testing as alternatives to the OGTT for diabetes screening. However, due to the limited sensitivity of each test when used alone, there is a risk of missed diagnoses, potentially missing a significant number of Pre-DM and T2D cases^12–14^. A research of 2,332 Chinese people discovered that the sensitivity of screening for Pre-DM using FPG was only 48.3%, indicating a significant percentage of missed diagnoses at 51.7%^15^. Another study found that using the HbA1c test alone missed (or underdiagnosed) 59% of patients with Pre-DM^16^. An accurate and early diagnosis of Pre-DM and T2D is a primary prerequisite for its effective prevention, control, and treatment^17^. Therefore, it is imperative to develop a practical and concise biomarker panel to identify individuals with Pre-DM and early T2D, thereby providing a more reliable diagnostic tool for large-scale screening and public health prevention and control strategies.

Metabolomics, which involves the systematic examination of dynamic changes in endogenous metabolites^18^, has the potential to disclose disease causes, discover biomarkers, and evaluate therapy efficacy^19–22^, displaying enormous promise in diabetes research^4^. The mainstream strategies (untargeted, targeted, and the emerging pseudo-targeted approach) each offer distinct advantages: untargeted metabolomics provides a broad-spectrum perspective, targeted enables precise quantification, while the pseudo-targeted method (e.g., Multiple Reaction Monitoring) combines wide coverage with high precision. These methods effectively identify lipid signatures specific to Pre-DM and T2D, for example, diacylglycerol, triacylglycerol (TG), phosphatidylcholine, significantly enhancing early diagnostic accuracy and reducing the risk of missed diagnoses^18,23–26^. Being an essential subfield of metabolomics, lipidomics offers crucial insights into the disturbances of lipid metabolism associated with diabetes^27–30^. Multiple prospective studies have confirmed that various metabolites, such as amino acids, glycerophospholipids (GPL), and purine nucleotides, undergo significant alterations before T2D onset^31–34^. These findings not only elucidate the metabolic abnormalities characteristic of the pre-disease stage but also provide crucial scientific rationale and potential biomarkers for early diagnosis, risk stratification, and personalized prevention.

Although the application of metabolomics in clinical research on Pre-DM and T2D has garnered increasing attention in recent years, there remains a lack of systematic reviews that comprehensively analyze current research trends, hotspots, and thematic evolution in this field. Bibliometrics, as a significant branch of information science, enables the quantitative assessment of literature quality, output volume, and citation impact within a specific research domain through statistical methods^35–37^. By employing retrospective data analysis, relationship mapping, and developmental forecasting, it has become an essential tool for identifying research trends, evaluating scholarly contributions, and predicting future directions^38,39^. Specifically, this methodology enables the extraction and integration of comprehensive data, including countries, institutions, journals, authors, references, and keywords, to evaluate the current landscape, emerging trends, and research frontiers within a given field. It further helps reveal academic collaboration networks, research hotspots, and evolving thematic directions^40^. In this study, we integrate the probabilistic modeling-based normalization approach of VOSviewer with the set theory-based normalization method of CiteSpace. Through collaboration network analysis, co-word analysis, clustering, and burst detection, we examine the evolutionary trajectory of research on metabolomics in Pre-DM and T2D across temporal and spatial dimensions. This provides a systematic and comprehensive presentation of the field’s cutting-edge hotspots and evolutionary trends.

## 2 Methods

### 2.1 Data Sources

All of the literature used in this study was obtained from the Web of Science Core Collection (WOSCC). To ensure the accuracy and quality of retrieval data, the Science Citation Index Expanded (SCI-E) and Social Sciences Citation Index (SSCI) databases were used. During the search, subject headings, free-text terms, and keywords were all employed simultaneously. Figure 1 illustrates the search approach, with the search scope spanning from the database’s inception to May 2025. To eliminate any bias from database upgrades, the data-gathering deadline was set for May 23, 2025.

**Figure 1.**
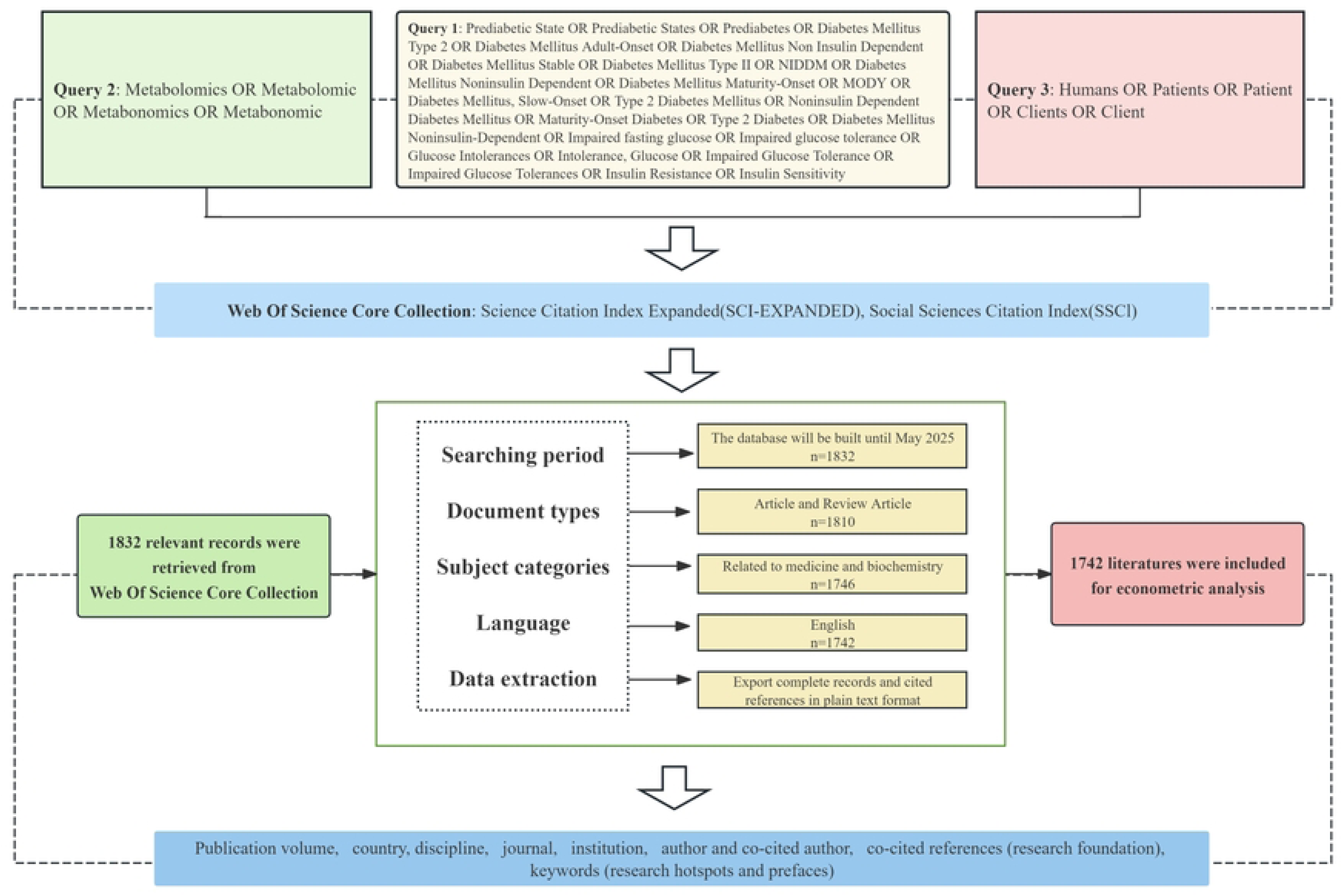
Flow Diagram of the Study Selection Process.

After a preliminary search, a total of 1,832 articles were retrieved, covering eight types of literature. Subsequently, they were screened, and the literature categories other than articles and review articles were excluded. The remaining articles numbered 1,810. Then, based on the discipline categories (based on Web of Science Categories with manual refinement), 1,746 articles related to medical-related disciplines were screened. Finally, manual verification was conducted, and four duplicate, non-English, and irrelevant articles related to this study were removed. Ultimately, this study comprised 1,742 articles from 75 countries and 2,937 institutions, authored by 13,505 researchers and published in 554 journals. These publications had 77,617 references from 7,649 different periodicals.

### 2.2 Data Analysis

VOSviewer (version 1.6.20) is a free Java-based freeware developed in 2009 by Van Eck and Waltman at Leiden University’s Centre for Science and Technology Studies. Due to its powerful graphical capabilities and ability to process large amounts of data^41^, the tool is extensively used to extract crucial information from voluminous publications, as well as to generate collaboration, co-citation, and co-occurrence networks^42,43^. CiteSpace (version 6.2.R3) is a bibliometric analysis and visualization software developed by Professor Chao Mei Chen of Drexel University in the United States^42,44^. It can visually present the foundational knowledge and research hotspots within the relevant field while also helping to predict its research frontiers^45^. The emergence, application, and development of these two software packages have accelerated research in the field of information visualization^38^. This study will utilize two software platforms to conduct statistical assessments of the research foundations, the current state, significant research areas, and emerging trends in the field of metabolomics for Pre-DM and T2D. The goal is to provide new insights and directions for the future growth of this study area.

## 3 Results

### 3.1 Annual Publication Analysis

The volume of literature published over a period reflects the current state and trends in research within a field. Through retrieval, 1,742 studies on metabolomics related to Pre-DM and T2D were identified, comprising 1,493 articles and 249 review articles. As shown in Figure 2, from 2003 to 2025, the number of publications related to metabolomics research in the Pre-DM and T2D field has exhibited an overall upward trend. Since 2013, this growth has remained largely stable. After 2018, annual publications on this topic routinely exceeded 100 papers, indicating that research in this area is in a relatively stable and continuously rising phase.

**Figure 2.**
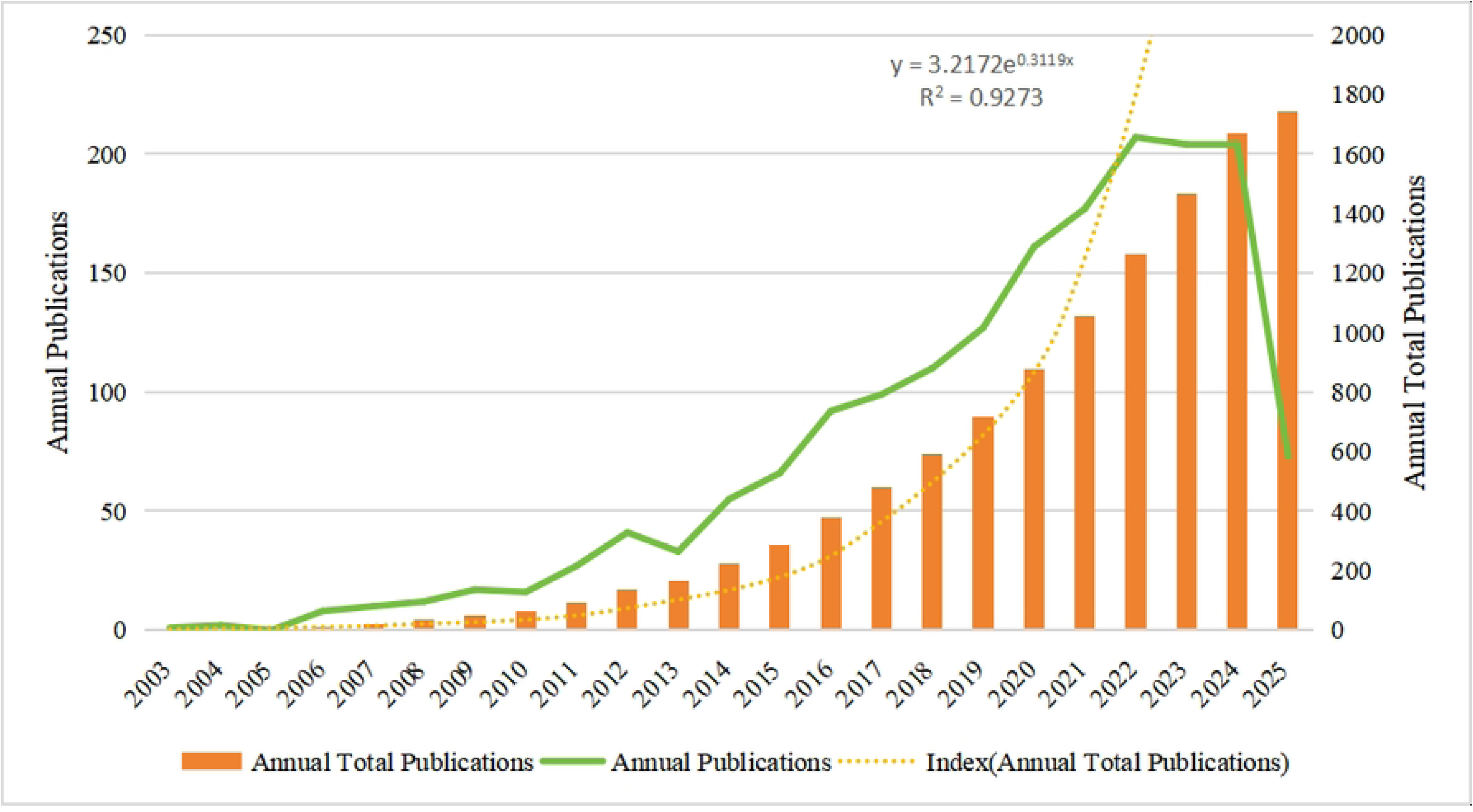
Distribution of Publications by Year

### 3.2 Country/Region Distribution

This study included articles from 75 countries and regions. The top 10 contributors predominantly came from Europe, North America, and Asia, with Europe (n = 7) accounting for the largest proportion (Table 1; Figure 3A). Among the top ten countries, China ranked first with 572 publications (32.84%), followed by the United States with 467 publications (26.81%). Germany placed third with 171 publications (9.82%), and England ranked fourth with 151 publications (8.67%). Overall, more than half of the publications came from China and the United States. However, in terms of total citations, the United States leads with 29,877, followed by China with 15,618, and the United Kingdom with 10,524. Although China publishes the most articles, the United States receives more citations than China. While England publishes fewer publications than China and the United States, the average number of citations per manuscript is higher.

**Figure 3.**
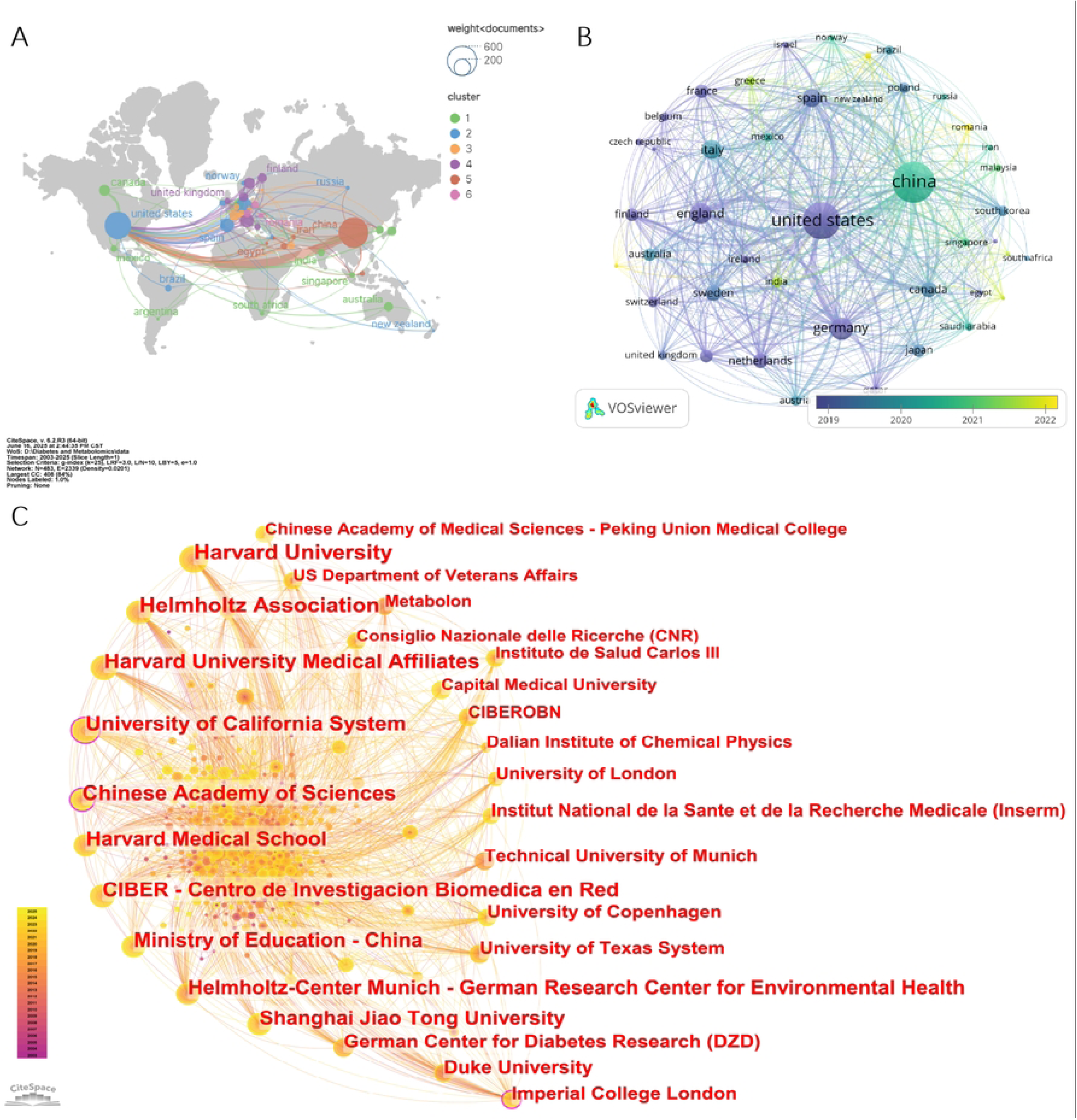
Bibliometric Analysis of Countries and Institutions in Metabolomics and Pre-DM/T2D Research. (A) Network map of international collaboration. (B) Cluster visualization of the country collaboration network. (C) Co-occurrence map of institutional collaborations.

**Table 1.** Top 10 Countries/Regions by Number of Publications.

| Rank | Country/region | Documents | Citations | Average citations | Proportion (%) |
| --- | --- | --- | --- | --- | --- |
| 1 | China (Asia) | 572 | 15618 | 27.30 | 32.84 |
| 2 | United States (North America) | 467 | 29877 | 63.98 | 26.81 |
| 3 | Germany (Europe) | 171 | 7867 | 46.01 | 9.82 |
| 4 | England (Europe) | 151 | 10524 | 69.70 | 8.67 |
| 5 | Italy (Europe) | 118 | 4711 | 39.92 | 6.77 |
| 6 | Spain (Europe) | 112 | 5051 | 45.10 | 6.43 |
| 7 | Canada (North America) | 79 | 3777 | 47.81 | 4.54 |
| 8 | Sweden (Europe) | 79 | 4350 | 55.06 | 4.54 |
| 9 | Netherlands (Europe) | 73 | 5288 | 72.44 | 4.19 |
| 10 | France (Europe) | 60 | 4226 | 70.43 | 3.44 |

A visualization analysis was conducted on the 44 countries among the 75 included in the study that had ≥5 publications. A collaborative network diagram was created using each country’s publishing volume and relationships, with node size representing publication volume and the thickness of the lines linking nodes reflecting the strength of linkages between nations (Figure 3B). It is evident that international collaboration in this field is extensive. For example, the United States has developed strong cooperative relationships with China, Canada, and Spain, while China has forged strong partnerships with the United States, Spain, Canada, Germany, and several other countries. Notably, academic collaboration between China and the United States is the most extensive. After superimposing the time dimension, the gradual change in colors can visually reflect the research situation of various countries in this field over recent years, with node colors indicating the temporal distribution (average year of paper publication). Figure 3B illustrates how countries such as Romania, Greece, and India have increasingly focused on this area of research in recent years.

### 3.3 Disciplines and Research Institutions

By summarizing the publication volume across relevant disciplines, it was found that the top three disciplines in terms of publication volume were Endocrinology and Metabolism (24.11%), Biochemistry and Molecular Biology (16.88%), and Nutrition and Dietetics (8.21%) (Table 2). Other fields included pharmacology and pharmacy (7.41%), multidisciplinary sciences (7.23%), experimental medicine research (7.00%), and biochemical research methods (5.68%).

**Table 2.**
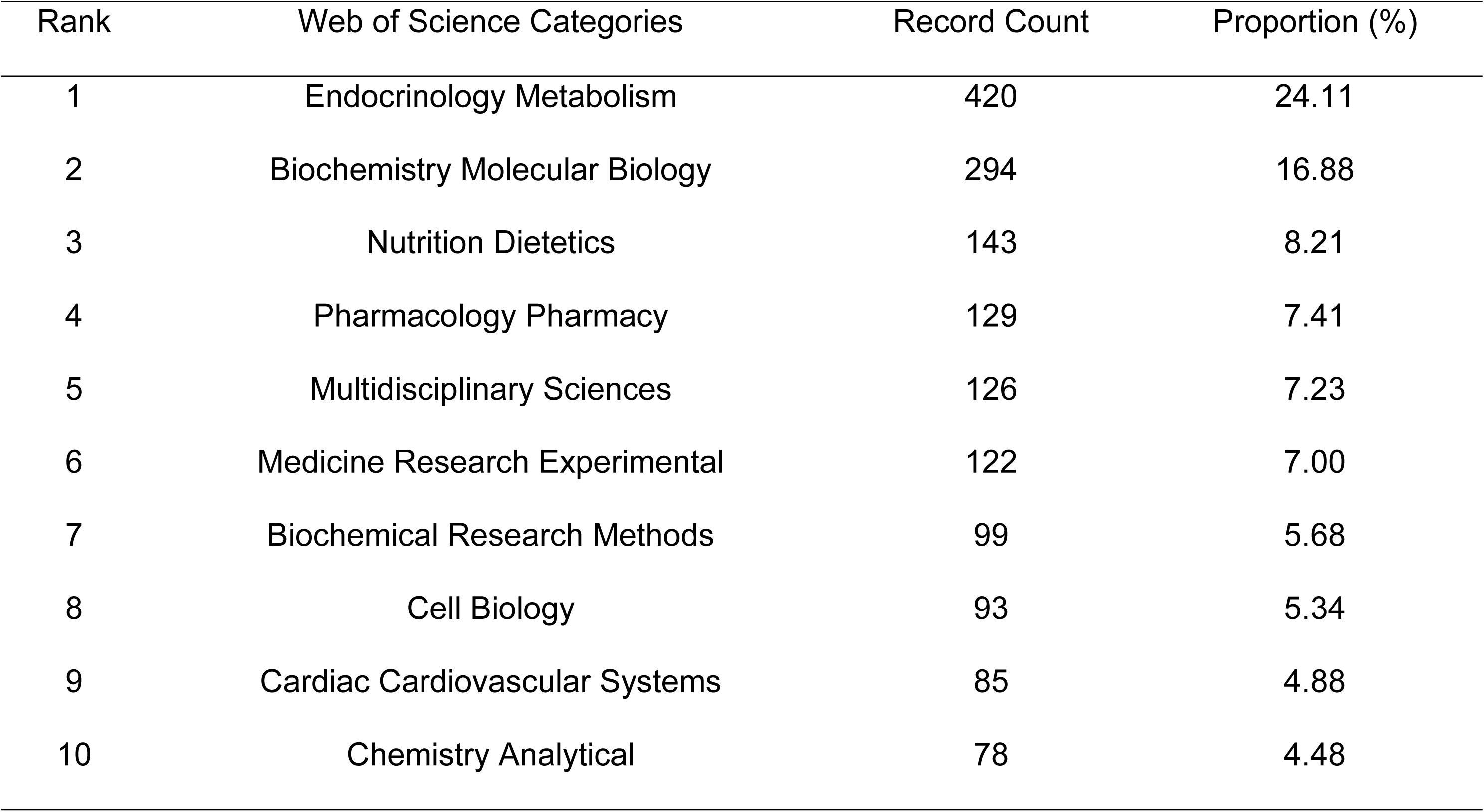
Top 10 Subject Areas in Metabolomics and Pre-DM/T2D Research.

Globally, Harvard University (77, 4.42%) and the Helmholtz Association (73, 4.19%) lead in paper output in this subject, with publications receiving 6,142 and 4,445 citations, respectively (Table 3). The top ten institutions include four from the United States, three from China, and two from Germany. Another is Spain’s Centro de Investigación Biomédica en Red (CIBER). The CiteSpace parameters were configured as follows: time slice (2003-2025), number of years per slice (1), node type (institution), and selection criteria (K = 25). Other parameters remained at their default values. A network co-occurrence graph with 483 nodes, 2339 connections, and 0.0201 density was generated (Figure 3C). The size of the nodes in the graph indicates the number of publications by the relevant institutions. The connecting lines between the nodes represent collaboration between the institutions. The node color, which ranges from dark to light, indicates the institutions’ activity in this research area since 2003. The number of connected lines indicates the degree of cooperation between each institution.

**Table 3.** Top 10 Most Productive Institutions in Metabolomics and Pre-DM/T2D Research.

| Rank | Institution | Country | Documents | Proportion<br>(%) | citations | Average<br>Citations | H-Index | Centrality | In the year |
| --- | --- | --- | --- | --- | --- | --- | --- | --- | --- |
| 1 | Harvard University | United States | 77 | 4.42 | 6142 | 79.77 | 36 | 0.05 | 2009-2025 |
| 2 | Helmholtz Association | Germany | 73 | 4.19 | 4445 | 60.89 | 32 | 0.09 | 2008-2025 |
| 3 | University of California System | United States | 66 | 3.79 | 3147 | 47.68 | 29 | 0.15 | 2010-2025 |
| 4 | Harvard University Medical Affiliates | United States | 65 | 3.73 | 5269 | 81.06 | 32 | 0.03 | 2009-2025 |
| 5 | Chinese Academy of Sciences | China | 60 | 3.44 | 3762 | 62.7 | 29 | 0.18 | 2004-2025 |
| 6 | Harvard Medical School | United States | 60 | 3.44 | 5527 | 92.12 | 30 | 0.06 | 2009-2025 |
| 7 | Shanghai Jiao Tong University | China | 58 | 3.33 | 2599 | 44.81 | 26 | 0.06 | 2009-2025 |
| 8 | Centro de Investigación Biomédica en Red | Spain | 57 | 3.27 | 3480 | 61.05 | 24 | 0.06 | 2012-2024 |
| 9 | Helmholtz Center Munich German Research Center for Environmental Health | Germany | 55 | 3.16 | 3708 | 67.42 | 28 | 0.09 | 2008-2024 |
| 10 | Ministry of Education China | China | 54 | 3.10 | 2857 | 52.91 | 23 | 0.06 | 2011-2025 |

### 3.4 Journals and Co-cited Journals

The 1,742 papers in metabolomics related to Pre-DM and T2D were published in 554 journals, of which 88 journals had ≥5 publications. Among the top ten journals (Table 4), *Metabolites* had the most articles (IF=3.5, Q2), followed by *Scientific Reports* (IF=3.8, Q1), *Journal of Proteome Research* (IF=3.8, Q1), and *Metabolomics* (IF=3.5, Q2). The journals with the highest impact factors were *Diabetologia* (IF = 8.4, Q1) and the Journal of Clinical Endocrinology and Metabolism (IF = 5.3, Q1). Subsequently, the aforementioned 88 journals were filtered using VOSviewer. As shown in Figure 4A, a strong co-citation link was observed between Metabolites and Clinical Nutrition, Diabetes Research and Clinical Practice, and other journals.

**Figure 4.**
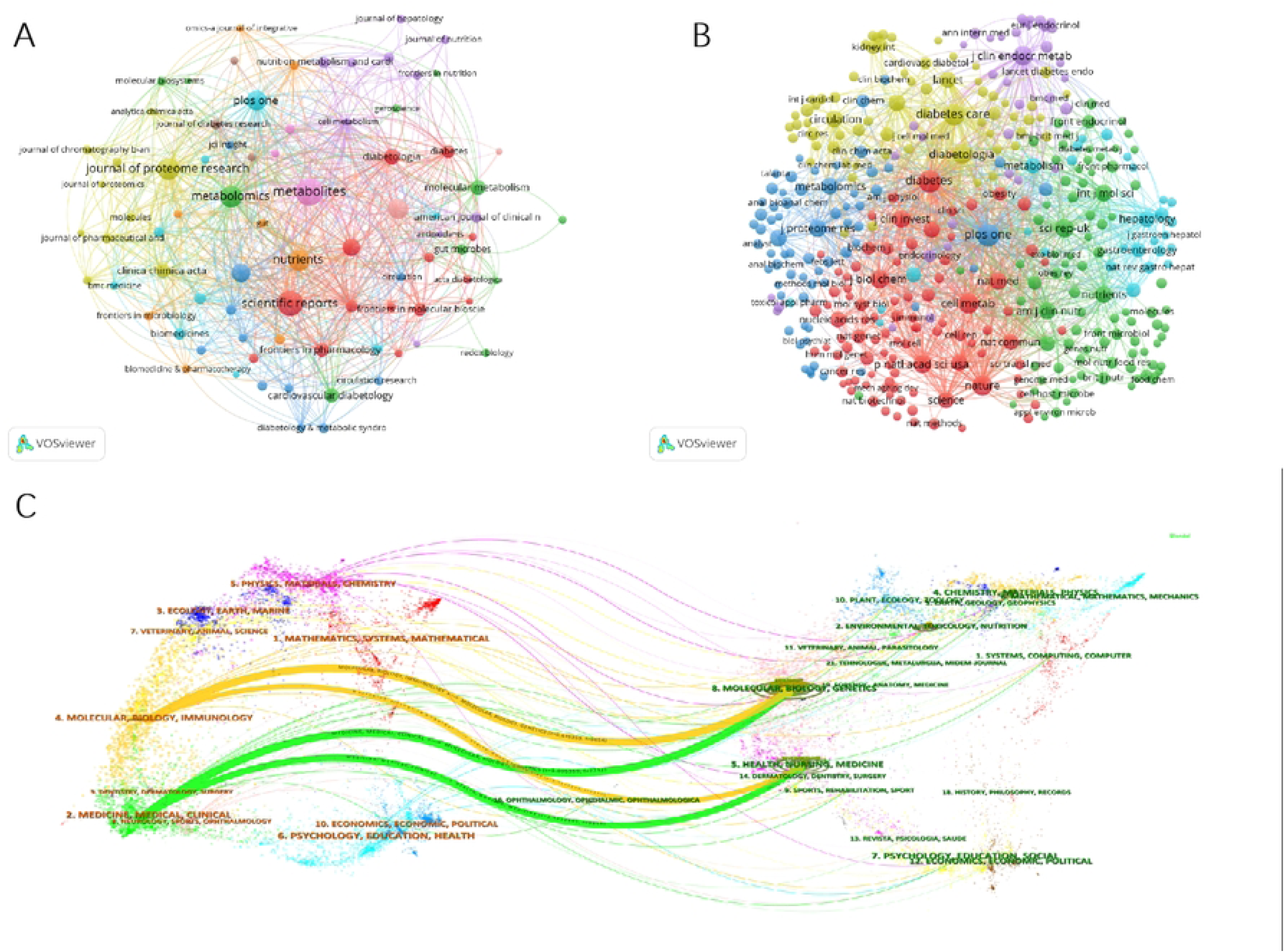
Journal Analysis in Metabolomics and Pre-DM/T2D Research. (A) Cluster visualization of the journal network. (B) Cluster analysis of the co-citation network. (C) Dual-map overlay of journals. Note: On the left are clusters of citing journals, which can represent the frontiers of knowledge in the relevant research area. On the right are clusters of cited journals, which represent the research base in the relevant research field. Individual labels represent the journals’ disciplinary divisions, whereas colored routes represent citation links between the left and right sides.

**Table 4.**
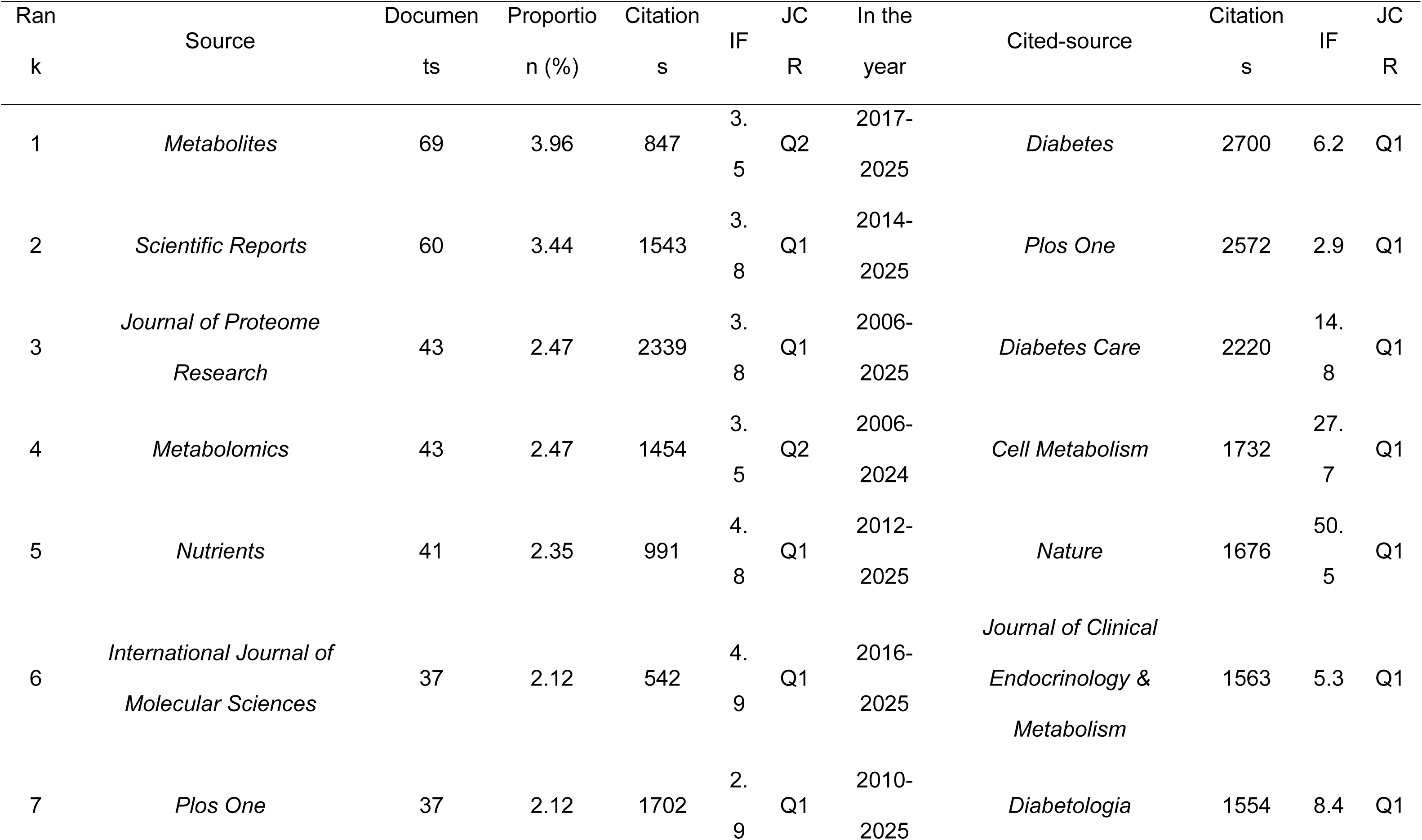
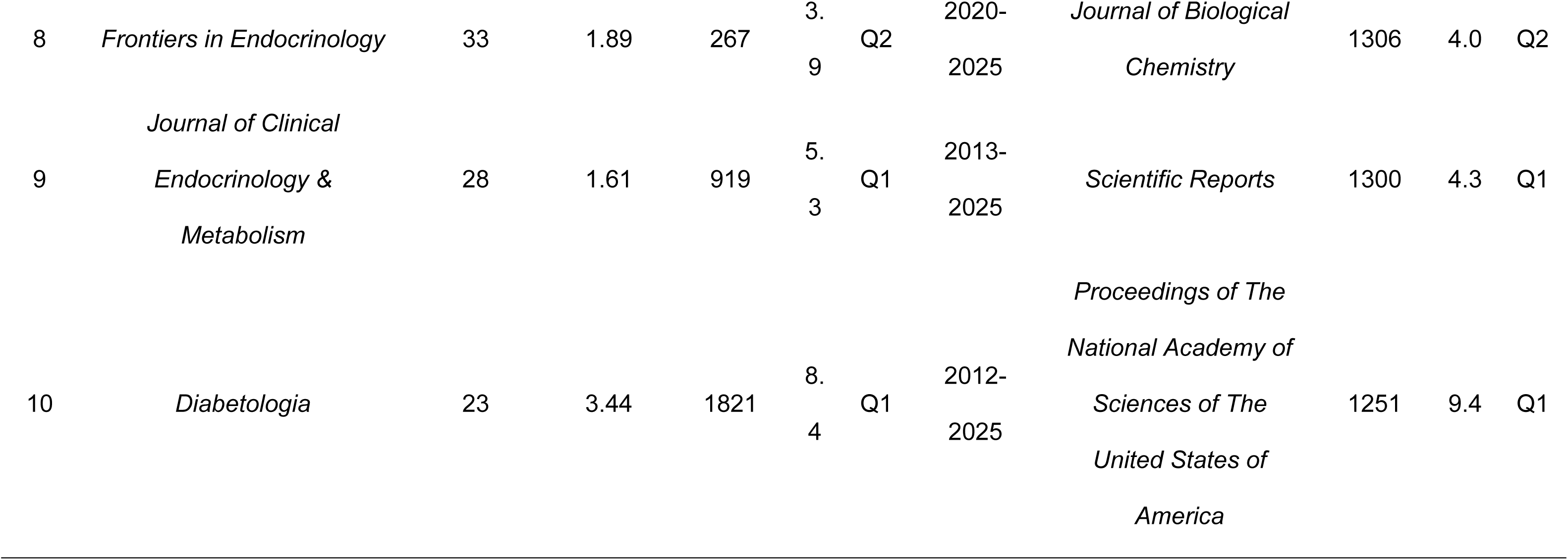
Top 10 Journals and Co-cited Journals in Metabolomics and Pre-DM/T2D Research.

| Rank | Source | Documents | Proportion (%) | Citations | IF | JCR | In the year | Cited-source | Citations | IF | JCR |
| --- | --- | --- | --- | --- | --- | --- | --- | --- | --- | --- | --- |
| 1 | *Metabolites* | 69 | 3.96 | 847 | 3.5 | Q2 | 2017-2025 | *Diabetes* | 2700 | 6.2 | Q1 |
| 2 | *Scientific Reports* | 60 | 3.44 | 1543 | 3.8 | Q1 | 2014-2025 | *Plos One* | 2572 | 2.9 | Q1 |
| 3 | *Journal of Proteome Research* | 43 | 2.47 | 2339 | 3.8 | Q1 | 2006-2025 | *Diabetes Care* | 2220 | 14.8 | Q1 |
| 4 | *Metabolomics* | 43 | 2.47 | 1454 | 3.5 | Q2 | 2006-2024 | *Cell Metabolism* | 1732 | 27.7 | Q1 |
| 5 | *Nutrients* | 41 | 2.35 | 991 | 4.8 | Q1 | 2012-2025 | *Nature* | 1676 | 50.5 | Q1 |
| 6 | *International Journal of Molecular Sciences* | 37 | 2.12 | 542 | 4.9 | Q1 | 2016-2025 | *Journal of Clinical Endocrinology & Metabolism* | 1563 | 5.3 | Q1 |
| 7 | *Plos One* | 37 | 2.12 | 1702 | 2.9 | Q1 | 2010-2025 | *Diabetologia* | 1554 | 8.4 | Q1 |
| 8 | *Frontiers in Endocrinology* | 33 | 1.89 | 267 | 3.<br>9 | Q2 | 2020-2025 | *Journal of Biological*<br>*Chemistry* | 1306 | 4.0 | Q2 |
| 9 | *Journal of Clinical*<br>*Endocrinology &*<br>*Metabolism* | 28 | 1.61 | 919 | 5.<br>3 | Q1 | 2013-2025 | *Scientific Reports* | 1300 | 4.3 | Q1 |
| 10 | *Diabetologia* | 23 | 3.44 | 1821 | 8.<br>4 | Q1 | 2012-2025 | *Proceedings of The*<br>*National Academy of*<br>*Sciences of The*<br>*United States of*<br>*America* | 1251 | 9.4 | Q1 |

Among the Top 10 most co-cited journals (Table 4), all journals had more than 1000 co-citations, with *Diabetes* (n=2700) being the most co-cited, followed by *Plos One* (n=2572) and *Diabetes Care* (n=2220). *Nature* (IF=50.5), *Cell Metabolism* (IF=27.7), and *Diabetes Care* (IF=14.8) have the highest impact factors of the top ten co-cited journals. VOSviewer was used to filter journals with a minimum co-citation of ≥ 50 to map the co-citation network. As shown in Figure 4B, all co-cited journals exhibited strong co-citation relationships with each other.

In Figure 4C, the dual graphical overlay of journals depicts the distribution of research topics as well as the citation relationships between journals and cited journals. We can see the four most important paths. Respectively represent that Molecular, Biology, Genetics journals and Health, Nursing, Medicine are frequently cited by Molecular/Biology/Immunology and Medicine/Medical/Clinical journals.

### 3.5 Author Analysis

This study included a total of 13,505 participants. The authors with the most publications are Jerzy Adamski, from Helmholtz Munich - German Research Center for Environmental Health (Germany), and Guowang Xu, from the Dalian Institute of Chemical Physics, Chinese Academy of Sciences (China). Four of the top ten authors have more than 20 publications (Table 5).

**Table 5.** Top 10 Most Productive Authors in Metabolomics and Pre-DM/T2D Research.

| Rank | Author | Documents | Citations | Average Citations | H-Index | Proportion(%) | In the year |
| --- | --- | --- | --- | --- | --- | --- | --- |
| 1 | Jerzy Adamski | 34 | 2227 | 65.5 | 18 | 1.95 | 2011-2024 |
| 2 | Guowang Xu | 27 | 1831 | 67.81 | 19 | 1.55 | 2004-2025 |
| 3 | Cornelia Prehn | 25 | 1847 | 73.88 | 15 | 1.44 | 2011-2024 |
| 4 | Christopher B. Newgard | 24 | 4926 | 205.25 | 20 | 1.38 | 2009-2022 |
| 5 | Gabi Kastenmuller | 17 | 1892 | 111.29 | 13 | 0.98 | 2011-2024 |
| 6 | Xinjie Zhao | 17 | 1014 | 59.65 | 15 | 0.98 | 2010-2022 |
| 7 | Olga Ilkayeva | 17 | 3961 | 233 | 15 | 0.98 | 2009-2022 |
| 8 | Weiping Jia | 17 | 778 | 45.76 | 13 | 0.98 | 2009-2022 |
| 9 | James R. Bain | 16 | 4294 | 268.38 | 15 | 0.92 | 2009-2021 |
| 10 | Robert D. Stevens | 15 | 4155 | 277 | 14 | 0.86 | 2009-2016 |

A collaborative network analysis of authors can provide information about the representative scholars and core research strengths in this research topic. After using VOSviewer software to assess the collaboration network of authors with ≥5 publications, 149 out of 13050 writers satisfied the requirement. VOSviewer categorized the above 149 authors into six clusters, as shown in Figure 5A, with tight collaborations between Jerzy Adamski, Gabi Kastenmueller, and Christian Gieger, as well as between Guowang Xu, Xinjie Zhao, and Weiping Jia, all of whom have close collaborations.

**Figure 5.**
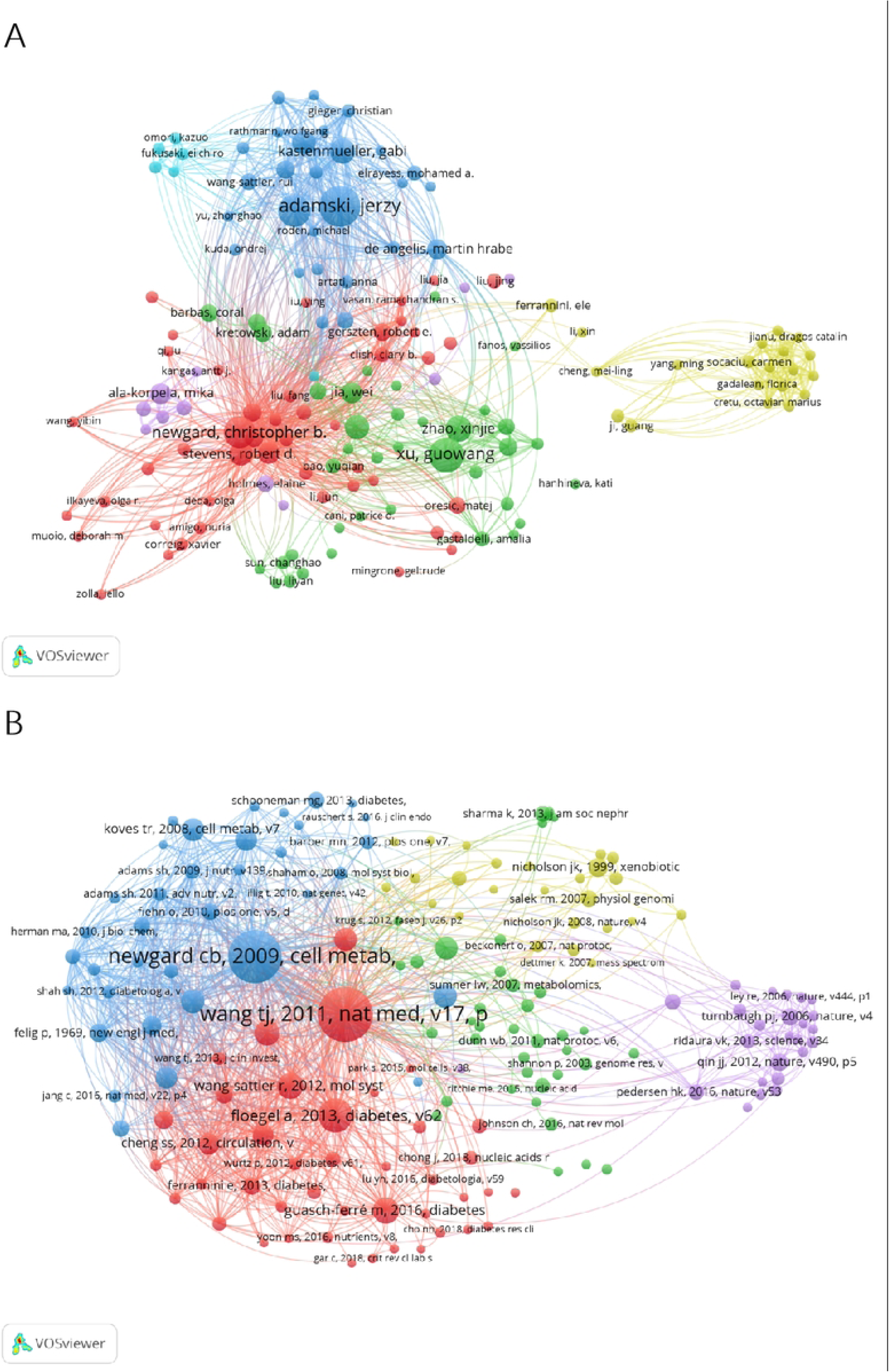
Analysis of Authors and Co-cited References in Metabolomics and Pre-DM/T2D Research. (A) Cluster visualization of the author collaboration network. (B) Cluster analysis of the reference co-citation network.

### 3.6 Reference Co-citation Analysis

Of the 77,617 co-cited references, the most cited article was “Metabolite profiles and the risk of developing diabetes” by Thomas J. Wang et al., published in the journal Nature Medicine in 2011, with a co-citation of 265, followed by “A branched-chain amino acid-related metabolic signature that differentiates obesity and lean humans and contributes to insulin resistance” by Christopher B. Newgard et al., published in the journal Anna Floegel in 2013, with a total of 251 citations (Table 6). A co-citation study of co-cited references using VOSviewer showed 166 publications with more than 20 citations. “Metabolite profiles and the risk of developing diabetes” remained central to this co-citation network (Figure 5B).

**Table 6.**
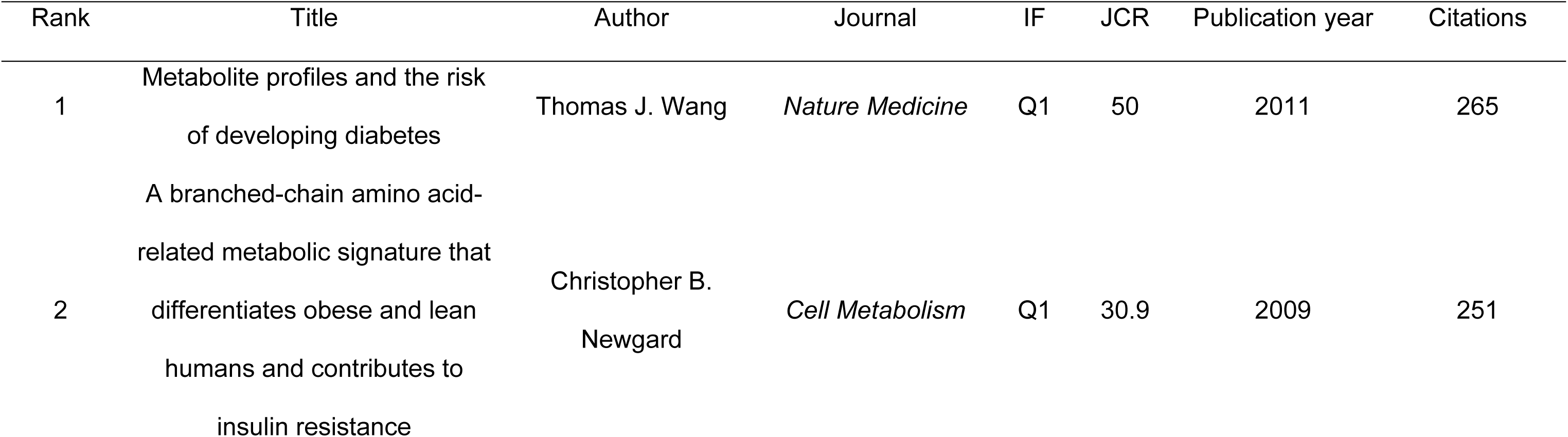
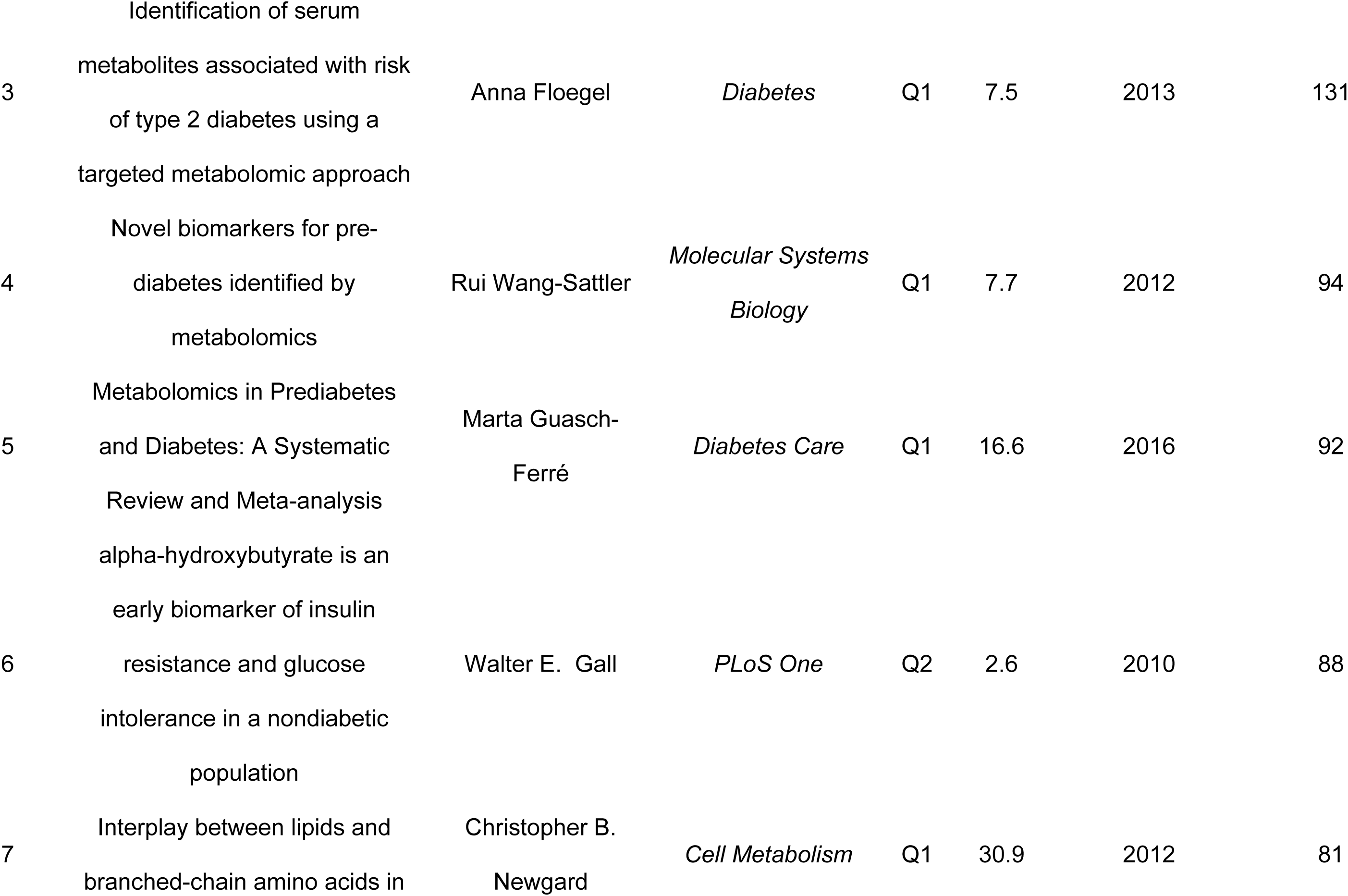
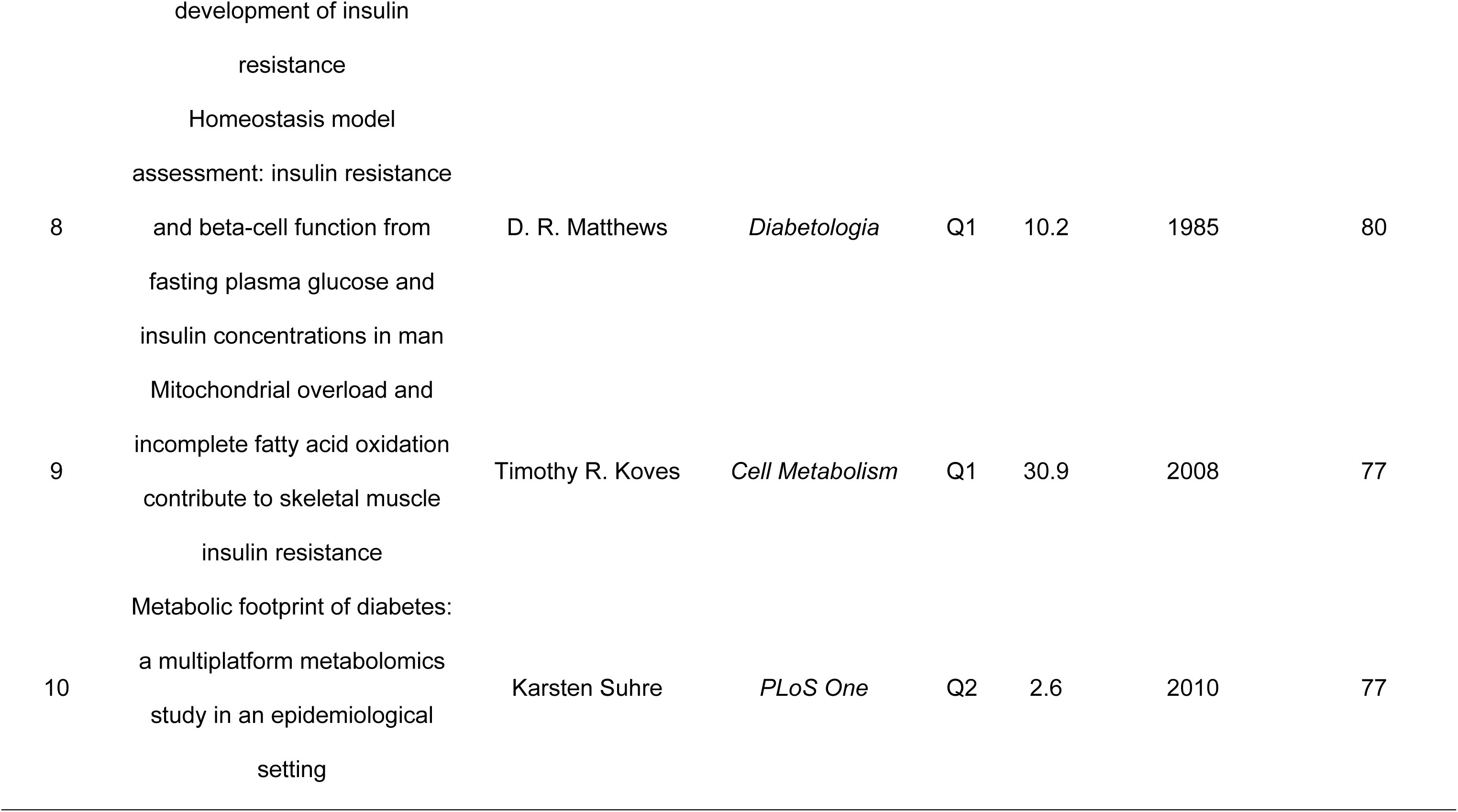
Top 10 Most Frequently Co-cited References in Metabolomics and Pre-DM/T2D Research.

### 3.7 Analysis of Research Hotspots and Frontiers

The titles and abstracts of the 1742 included articles contained a total of 6654 keywords. As shown in Table 7, the keywords “Metabolomics” (n=762), “Insulin-Resistance” (n=612), “Obesity” (n=318), “Risk” (n=266), “Biomarkers” (n=220), and “Metabolism” (n=215) all appeared more than 200 times, indicating that the research areas in this field of study are primarily focused on “Metabolomics” and “Insulin Resistance”.

**Table 7.** Top 20 Most Frequent Keywords in Metabolomics and Pre-DM/T2D Research.

| Rank | Keyword | Occurrences | Total link strength | Rank | Keyword | Occurrences | Total link strength |
| --- | --- | --- | --- | --- | --- | --- | --- |
| 1 | metabolomics | 762 | 1608 | 11 | inflammation | 133 | 313 |
| 2 | insulin-resistance | 612 | 1249 | 12 | association | 122 | 308 |
| 3 | obesity | 318 | 786 | 13 | diabetes | 119 | 266 |
| 4 | risk | 266 | 646 | 14 | oxidative stress | 117 | 258 |
| 5 | biomarkers | 220 | 558 | 15 | disease | 112 | 247 |
| 6 | metabolism | 215 | 438 | 16 | insulin resistance | 112 | 271 |
| 7 | type 2 diabetes | 147 | 338 | 17 | expression | 107 | 243 |
| 8 | glucose | 142 | 329 | 18 | metabolic syndrome | 104 | 277 |
| 9 | plasma | 141 | 371 | 19 | serum | 101 | 260 |
| 10 | gut microbiota | 137 | 293 | 20 | type 2 diabetes mellitus | 91 | 199 |

The VOSviewer software was used to analyse the co-occurrences of terms with ≥5 occurrences, yielding 668 keywords. As illustrated in Figure 6A, the size of the nodes indicated the frequency of keyword occurrence, with the closer the connection between the nodes, the greater the probability of two keywords occurring concurrently. These keywords were classified into five clusters, indicating the primary directions of metabolomics related to Pre-DM and T2D research. Cluster 1 (red) focuses on the application of metabolomics technologies, aiming to discover and validate biomarkers for early warning and risk assessment of diabetes. In contrast, Cluster 2 (green) is dedicated to the regulatory mechanisms of glucose metabolism, investigating fundamental metabolic processes at the molecular level, such as gene expression. Cluster 3 (blue) focuses on obesity, examining anthropometric indices such as body mass index (BMI) and their associations with diabetes, which are significant risk factors for the condition. Cluster 4 (yellow) delves into the specific metabolic basis of IR, with an emphasis on the role of amino acid metabolism disorders, including branched-chain amino acids (BCAAs) and acylcarnitines. Meanwhile, Cluster 5 (purple) highlights the central role of inflammation in driving diabetes-related metabolic diseases, linking immune responses to systemic metabolic dysregulation.

**Figure 6.**
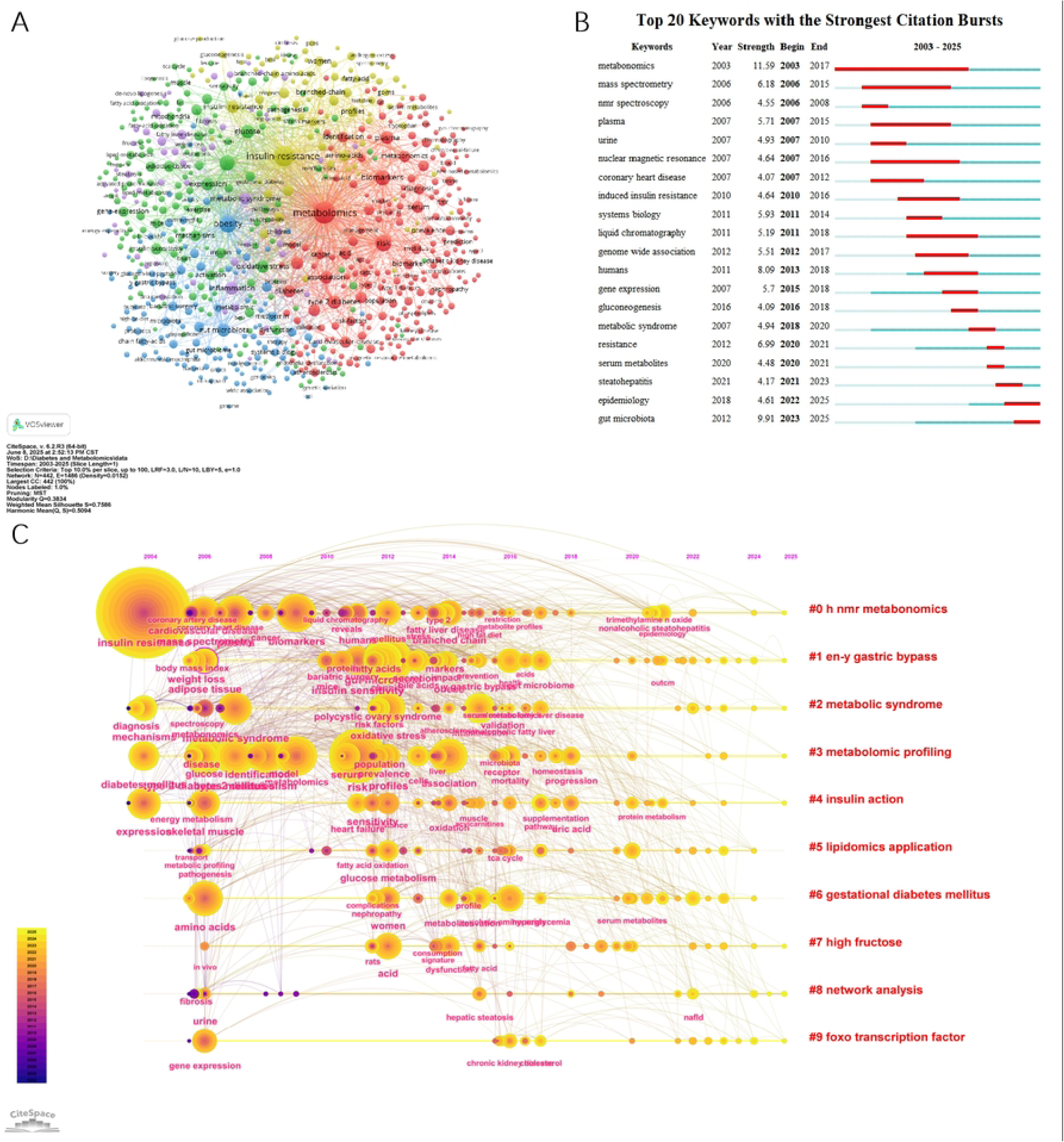
Keyword Analysis in Metabolomics and Pre-DM/T2D Research. (A) Cluster visualization of the keyword co-occurrence network. (B) Top 20 keywords with the strongest citation bursts (C) Timeline visualization of keyword clusters.

Overall, these five research clusters collectively form a multi-level and multi-faceted integrated research network centered on metabolomics and clinical diabetes research. All clusters focus on the field of diabetes health, with particular emphasis on the etiology, biomarker discovery, and pathophysiological mechanisms of obesity and its related complications. While each cluster remains relatively independent with its own distinct research focus, they are also closely interlinked and mutually informative, together revealing a comprehensive disease spectrum of diabetes ranging from molecular mechanisms to clinical phenotypes.

Keyword bursts refer to the terms that exhibit the highest frequency change over a certain period, which can reflect the issues of widespread interest among academics in the field. In this study, based on the keyword co-citation network, a burst detection analysis was conducted, identifying the top 20 keywords with the strongest citation bursts. As depicted in Figure 6B, the blue line represents the timeline, and the red paragraphs on the blue timeline represent burst detection, including the start year, finish year, and duration. Based on the time of appearance, “Metabonomics” (burst intensity 11.22), “Plasma” (burst intensity 6.74), and “Mass Spectrometry” (burst intensity 6.69), among others, were the primary focus of interest in the early days of the field. The primary study themes in the recent decade have been “Gut Microbiota” (burst intensity 8.82), “Humans” (burst intensity 8.01), “Resistance” (burst intensity 7.23), and “Metabolic Syndrome” (burst intensity 6.05).

Furthermore, the terms “Diabetic Kidney Disease” (burst intensity 5.43), “Epidemiology” (burst intensity 5.06), and “Gut Microbiota” (burst intensity 8.82) are on the rise and are at the forefront of study in their fields. The keyword timeline graph can display the dynamic evolution path of research hotspots represented by keywords, as well as investigate the temporal characteristics of research areas expressed through clustering and the rise and fall process of hot keyword research. According to the CiteSpace settings, a keyword timeline graph with 442 nodes, 1,486 connections, and a density of 0.0152 was generated (Fig. 6C), which visualizes the research hotspots and development directions of metabolomics related to Pre-DM and T2D in the temporal dimension. The image depicts nine clusters: ^1^H NMR Metabolomics, Roux-en-Y Gastric Bypass, Metabolic Syndrome, Metabolomic Profiling, Insulin Action, Lipidomics Applications, Gestational Diabetes Mellitus (GDM), High-Fructose Diet, Network Analysis, and the FOXO Transcription Factors. The picture shows that the majority of the research between 2004 and 2006 focused on IR, BMI, adipose tissue, spectroscopy, metabonomics, metabolic syndrome, glucose identification, energy metabolism, gene expression, metabolic profiling, amino acids, and gene expression. Between 2020 and 2022, the research mostly focused on trimethylamine n-oxide (TMAO), nonalcoholic steatohepatitis (NASH), epidemiology, protein metabolism, serum metabolites, and non-alcoholic fatty liver disease (NAFLD).

In summary, by integrating keyword burst detection and timeline analysis, this study clearly delineates a dynamic evolutionary panorama of metabolomics in the field of Pre-DM and T2D research, revealing a distinct three-phase developmental trajectory characterized by “technology-driven-mechanism exploration-clinical integration.” The early research focus was primarily technology-driven, centered on methodological advances such as mass spectrometry, metabolomics, and spectroscopy. The core objective was establishing analytical platforms and initially exploring associations with macro-phenotypes like IR, metabolic syndrome, and obesity. The mid-phase research emphasis shifted toward in-depth mechanistic investigation, focusing on specific pathways such as gut microbiota, lipidomics, and particular amino acid metabolism. This period revealed the inherent biological processes underlying metabolic disorders and expanded research into specific populations, including GDM. The current research priority has further evolved into clinical integration, emphasizing the precision prevention and management of emerging disease entities (e.g., NASH and diabetic kidney disease) and the accumulation of large-scale epidemiological evidence. There is a heightened focus on the role of host-microbiota co-metabolites (e.g., TMAO), marking the field’s progression toward clinical translation and deepened precision medicine. From a trend perspective, this field has undergone a spiral ascent from macro to micro and back to macro. This involves establishing methodologies, correlating macro-level phenotypes, delving into molecular mechanisms, and ultimately returning to address macro-level clinical problems, while now equipped with more powerful technical tools and deeper mechanistic insights. Looking forward, research on the gut microbiota, epidemiological big data, and metabolomics of specific complications will continue to serve as the core drivers advancing the discipline, ultimately contributing to early disease warning, stratified diagnosis, and personalized treatment.

## 4 Discussion

### 4.1 General Information

Based on the WOSCC database, we retrieved clinically oriented research publications related to metabolomics related to Pre-DM and T2D published up to May 2025 and conducted a bibliometric visualization analysis. This study provides a comprehensive overview of the current research landscape, identifies emerging trends, tracks disciplinary developments, and offers insights into future research directions. In recent years, the number of publications on the association between metabolomics related to Pre-DM and T2D has steadily increased, with a substantial peak in 2022, indicating the growing relevance of research in this area. This tendency may be related to the following factors: (1) The global prevalence of diabetes and other related metabolic diseases continues to rise^17^; (2) Metabolomics holds significant potential in diabetes research, effectively revealing key differential metabolites between diabetic patients and healthy individuals and identifying biomarkers associated with Pre-DM and T2D in subclinical states^46^, thus advancing early diagnosis and prevention of diabetes; (3) Advancements in metabolomic analytical technologies, coupled with the emergence of multi-omics integrated research paradigms^47^ and the widespread application and continuous innovation of bioinformatic methods—such as machine learning^48^—have enabled large-scale, high-throughput metabolite screening and identification.

China, the United States, and Germany have the highest number of articles on metabolomics related to Pre-DM and T2D. Articles from the United States and China have the highest citation frequency, indicating their significant academic influence. Furthermore, four of the top ten institutions by article output are in the United States, with three from China. This phenomenon may be explained by the high prevalence of diabetes-related metabolic illnesses in the United States and China, which has prompted the development of relevant laboratory and clinical research.

As summarized in this review, journals such as *Metabolites* and *Scientific Reports* serve as primary and popular platforms for publishing research in this field. In contrast, high-impact journals, including *Diabetes Care* and *Cell Metabolism*, constitute the core knowledge base for the field’s citation index. Researchers in this field should prioritize high-impact journals and publications to follow and improve the latest global research trends and developments. Such a suggestion will enhance the depth and breadth of the study, thereby increasing the academic influence and contributions to the field. From our analysis of the relationship between journals and cited journals, we also found that findings from fundamental disciplines, such as molecular biology and genetics, as well as applied literature in fields like health and clinical medicine, are frequently cited in immunology and clinical medical research. This reflects a deep interdisciplinary integration and contributions of basic research to clinical practice. Furthermore, such citation patterns suggest that traditional journal classifications fall short in fully capturing the multidisciplinary nature of research. Future academic undertakings should place greater emphasis on the multidimensional integration of knowledge flows.

From our analysis of author contributions, we found that Professor Jerzy Adamski of the Helmholtz Munich is the most prolific author in this bibliometric analysis. His team has systematically established the application framework of metabolomics in diabetes research, encompassing methodologies, biomarker discoveries, and mechanism studies of this disease^49^. Methodologically, the researchers clarified the distinguishing features of plasma and serum in metabolomic analysis^50^. In terms of biomarker discovery, the team identified elevated levels of BCAAs and reduced glycine (Gly) as robust predictive indicators for Pre-DM^12^. Additionally, they also discovered the significance of postprandial metabolites and novel N-acyl amino acids in the progression of disease^51,52^. Mechanistically, the team revealed that metformin modulates lipid metabolism via the adenosine monophosphate-activated protein kinase (AMPK) signaling pathway and influences citrulline metabolism through the AMPK-endothelial nitric oxide synthase (eNOS) signaling axis, providing a molecular basis for its cardiovascular protective effects^53,54^.

Another important contributor to this field, Professor Guowang Xu from the Dalian Institute of Chemical Physics, Chinese Academy of Sciences, has employed multidimensional metabolomics technologies to elucidate key metabolic features and underlying mechanisms associated with both diabetes and its Pre-DM states. On the methodological front, the team developed cutting-edge techniques, including pseudo-targeted two-dimensional liquid chromatography-mass spectrometry (2D LC-MS)^55^ and metabolome-exposome association study (mEWAS)^56^. They further integrated strategies, including lipidomics^7^, liquid chromatography-mass spectrometry (LC-MS)^57^, and comprehensive two-dimensional gas chromatography/time-of-flight mass spectrometry (GC×GC-TOFMS) ^58^, significantly enhancing metabolite detection capabilities. Regarding mechanisms and biomarkers, the team systematically elucidated key features of Pre-DM, such as elevated free fatty acids(FFAs) and dysregulated amino acid metabolism^17,55,57,59–61^. They identified that N-acetylhistamine drives disease progression by inhibiting AMPK phosphorylation^17^ and discovered susceptibility genes, including fatty acid desaturase 1 (FADS1) and WW domain-containing oxidoreductase (WWOX), through metabolome-genome-wide association studies (mGWAS)^62^. In terms of interventions, the team delineated the mechanisms by which berberine, exercise, and Roux-en-Y gastric bypass surgery exert therapeutic effects via the modulation of metabolic pathways^60,61,63^.

Professor Jerzy Adamski and Professor Guowang Xu are both global leaders in metabolomics-driven diabetes research, and their work demonstrates notable complementarity. Together, they have demonstrated the irreplaceable value of metabolomics in revealing early metabolic dysregulation in diabetes^23,57,59,64,65^, discovering novel biomarkers^61,65^, elucidating pathological mechanisms^17,60,66^, and evaluating therapeutic efficacy. Their contributions have collectively advanced the field toward more precise, systematic, and personalized approaches.

### 4.2 The Core Intellectual Base: A Co-citation Analysis

The summarization of the highly co-cited literature identified in this study elucidates not only the current research landscape and developmental trends of metabolomics in Pre-DM and T2D but also highlights that specific blood metabolites hold significant predictive value before the onset of diabetes.

The most cited study by Thomas J. Wang^67^ et al. (2011) employed mass spectrometry to elevate BCAA and aromatic amino acids (AAAs)levels. Especially, he discovered that the isoleucine(Ile), leucine (Leu), Valine (Val), Tyrosine (Tyr), and Phenylalanine (Phe) were significantly associated with Pre-DM, and subsequently developed a diabetes prediction model based on Ile, Tyr, and Phe. This finding is consistent with the second most co-cited publication, a 2009 study by Christopher B. Newgard^68^ et al. in *Cell Metabolism*, which reported that BCAA-related metabolites in obesity individuals are directly linked to IR, identifying it as an independent risk factor promoting the development of diabetes. Furthermore, the third co-citation written by Anna Floegel^69^ et al. (2013) in *Diabetes*, extended the spectrum of relevant metabolic markers to include sugar-derived metabolites and choline-containing phospholipids, confirming that disturbances across multiple metabolic pathways are independently associated with the risk of T2D. Notably, studies by Walter E. Gall^70^ et al. found that α-hydroxybutyrate (α-HB) can distinguish between individuals with IR and those with insulin sensitivity, as well as between those with standard glucose tolerance and those with IGT. Furthermore, it is independently associated with IR and impaired glucose regulation (IGR). Therefore, α-HB is considered an early biomarker for IR and IGR.

In summary, previous research has shown that metabolites such as carbohydrates (glucose and fructose), lipids (phospholipids, sphingomyelin, and TG), and amino acids (BCAAs, AAAs, Gly, and glutamine) have strong potential correlations with Pre-DM and T2D^71–74^. In the general population, these metabolic indicators have previously been shown to have the ability to diagnose diabetes and its associated complications^46^.

Collectively, these foundational studies establish a core conclusion: blood metabolic profiles can not only reveal the subclinical stage of diabetes and elucidate its pathogenic mechanisms but also ultimately provide a crucial scientific basis for early warning, risk stratification, and targeted personalized intervention of the disease.

### 4.3 Hotspots and Frontiers

Based on keyword co-occurrence, we can identify key themes and trends in metabolomics-related clinical research concerning Pre-DM and T2D over the past two decades. These core keywords include metabolomics, IR, obesity, risk, biomarkers, gut microbiota, amino acids, lipidomics, and mass spectrometry, among others.

#### 4.3.1 The Central Role of Insulin Resistance

IR serves as a common pathological basis for a range of highly prevalent diseases, including T2D, obesity, and cardiovascular diseases^75^. It is characterized by impaired cellular capacity for the storage and oxidation of glucose and lipids^66^, often accompanied by “metabolic inflexibility”, which is the inability to adapt oxidative pathways efficiently in response to fuel availability^76^. IR can lead to reduced glucose oxidation and insufficient suppression of lipid breakdown under insulin stimulation, resulting in ectopic fat accumulation and lipotoxicity, which in turn further exacerbates IR^77^, which is highly prevalent in obesity individuals, accounting for more than 80% of all T2D cases^61^.

The Pre-DM stage, characterized by IFG and/or IGT, represents a state of dysglycemia. Its core pathophysiological basis lies in the presence of IR in muscle, liver, and adipose tissue, coupled with early-phase insulin secretory dysfunction^57^. With the combined effects of insulin secretory deficiency and IR, β-cell function gradually decompensates, ultimately leading to the manifestation of overt hyperglycemia characteristic of T2D^51^. Upon progression to the overt T2D stage, patients may then face a significantly elevated risk of developing complications, including myocardial infarction, stroke, neuropathy, retinopathy, and renal failure^51^. Recent studies reported that carbohydrates (e.g., lactate and mannose), as well as lysophosphatidylcholines (LPC) and acylcarnitines, are associated with IR and IGT, beyond the normal glucose index^4^.

In summary, IR acts as a central driving factor throughout the entire disease continuum, from obesity and Pre-DM to overt T2D and its complications. Metabolomic studies are pivotal in elucidating the distinct metabolic signatures driven by IR, highlighting the fundamental importance of targeting IR for early intervention to halt disease progression and improve patient outcomes.

#### 4.3.2 Pre-DM and T2D-Associated Metabolic Biomarkers

From our bibliometric analysis, we filtered and summarized that the alterations in blood glucose levels, lipid metabolites (e.g., GPL, FFAs), amino acid-related metabolites (e.g., BCAAs, AAAs), as well as gut microbiota-derived metabolites (e.g., hippuric acid [HA] and methylxanthines [MXs]), collectively form a characteristic network of metabolic disturbances in both Pre-DM and T2D. A brief summary of the metabolomic analytical workflow and characteristic differential metabolites associated with diabetes is provided in Figure 7.

**Figure 7.**
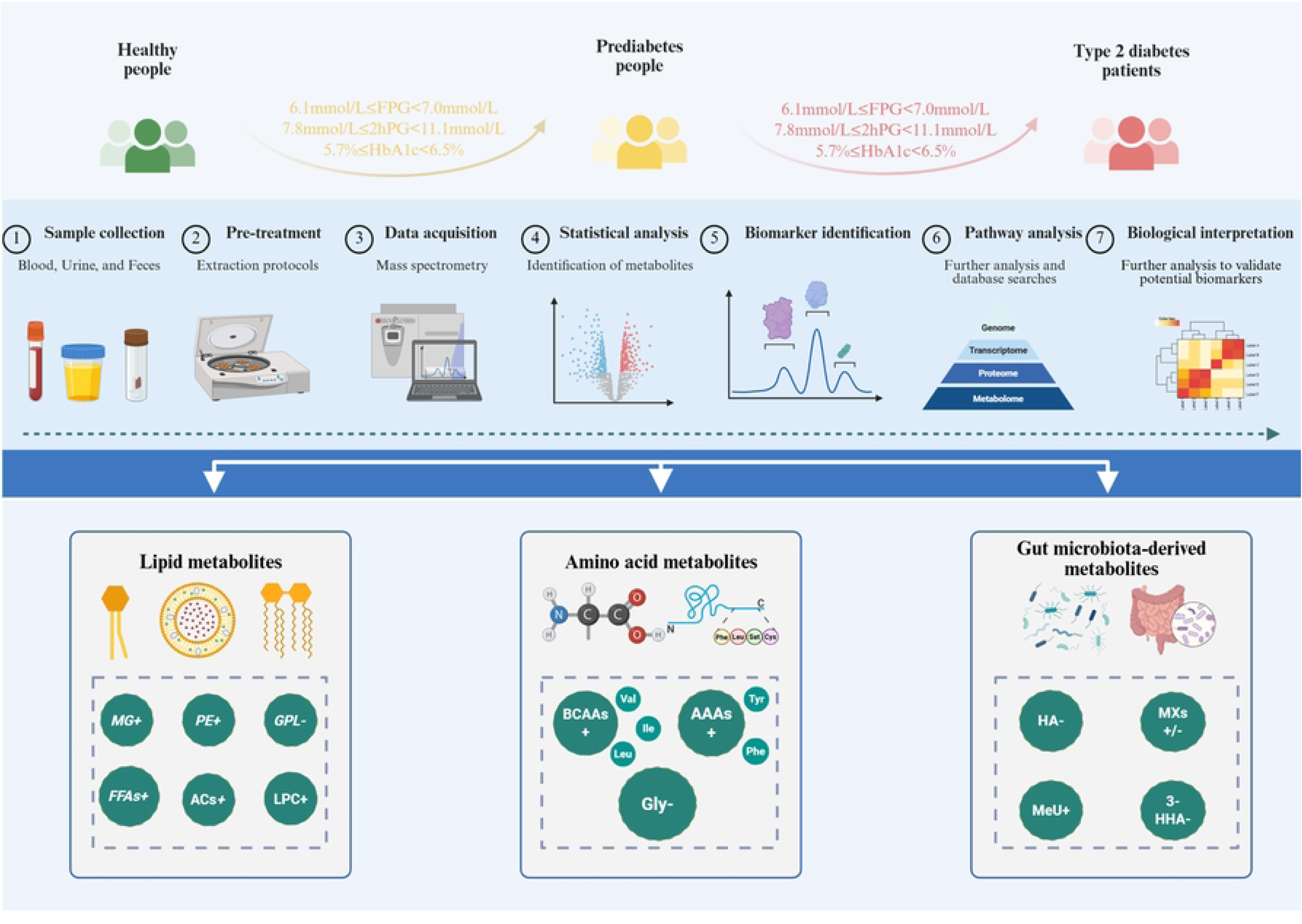
Metabolic Workflow and Characteristic Metabolites in Pre-DM/T2D.

##### 4.3.2.1 Lipid Metabolites

Individuals with Pre-DM already exhibit characteristic alterations in their lipid profiles. These abnormal lipid signatures can serve as early indicators of metabolic dysregulation and predict the risk of progression to overt T2D ^78^. Beyond traditional lipid markers extensively linked to diabetes^79,80^, such as plasma TG, total cholesterol (TC), low-density lipoprotein cholesterol (LDL-C), and high-density lipoprotein cholesterol (HDL-C). Recent studies have identified multiple specific lipid molecules closely associated with the onset and progression of disease. For instance, abnormalities in lipid metabolites such as monoglycerides (MG), phosphatidylethanolamine (PE) ^81^, GPL, FFAs, acylcarnitines (ACs), and LPC^82,32,83^ are not only associated with metabolic dysregulation in individuals with impaired glucose tolerance but also contribute to disease progression^71^. Notably, elevated levels of multiple fatty acids (FAs) are observed in diabetic patients^55^. These elevated FAs can induce IR and β-cell dysfunction, thereby exacerbating the disease^55,84,85^. During the progression from Pre-DM to T2D, lipid metabolism disorders such as ceramide, PE, diacylglycerol (DG), and TG progressively worsened^23^. This not only reveals the potential value of FAs receptors as therapeutic targets but also underscores the central role of abnormal lipid metabolism in the onset and development of diabetes. Therefore, lipidomics analysis overcomes the limitations of traditional lipid testing, providing a high-resolution perspective on the dynamic changes as well as more detailed detection of subclassified molecules in lipid metabolism during the early progression of diabetes and opening new dimensions for understanding disease mechanisms.

##### 4.3.2.2 Amino Acid Metabolites

Disorders of amino acid metabolism play a significant role in the development and progression of T2D. This is primarily characterized by abnormally elevated plasma levels of BCAAs and AAAs^86^, which are significantly associated with insulin IR, β-cell dysfunction, and an increased risk of diabetes^67^. Notably, elevated concentrations of BCAAs can predict disease onset up to a decade in advance^87^. Conversely, Gly, a product associated with high insulin levels^71^, demonstrates an inverse correlation with diabetes risk^88^. Recent studies have identified novel metabolites, such as picolinoylglycine and N-lactoyl-amino acids, as being involved in the transition from Pre-DM to overt T2D^51^. In summary, amino acids and their derivatives play fundamental roles not only in the pathogenesis and progression of T2D but also serve as crucial biomarkers for early diagnosis and risk prediction. Abnormal changes in traditional amino acids can reflect IR and β-cell dysfunction earlier than conventional indicators, providing crucial evidence for screening and intervention in Pre-DM and T2D. The discovery of novel amino acid-derived metabolites further deepens our understanding of the progression mechanisms of diabetes, opening new avenues for developing more precise early warning strategies.

##### 4.3.2.3 Gut Microbiota-derived Metabolites

From our analysis, we also found that recently, more and more metabolomic studies have been combined with gut microbiota analysis. Disrupted gut microbiota contributes to the pathogenesis of Pre-DM and T2D by regulating multiple metabolic pathways^89^. Specific mechanisms include modulating the metabolism of BCAAs and AAAs^90,91^, as well as altering the metabolism of short-chain fatty acids (SCFAs)^92^ and bile acids^89^, thereby interfering with host glucose and lipid homeostasis. Studies have shown that levels of microbiota-derived metabolites, such as imidazole propionate, trimethylamine N-oxide (TMAO), and succinate, are altered in patients with T2D. These metabolites can impair insulin signaling and glucose tolerance through pathways including the mechanistic target of rapamycin complex 1 (mTORC1) and AMPK^17,91,93,94^. Another large-scale metabolomics study further revealed alterations in 111 microbiota-associated metabolites in the serum of T2D patients, with 103 of these significantly correlated with FBG and HbA1c. These alterations primarily involved metabolic pathways of BCAAs, AAAs, and unsaturated fatty acids (UFAs)^17^. Furthermore, Pre-DM individuals already exhibit abnormal levels of microbiota metabolites such as HA, MXs, methyluric acids (MeU), and 3-hydroxyhippuric acid (3-HHA)^57^, suggesting that gut microbiota metabolic dysregulation precedes the clinical onset of T2D. These findings offer unique insights into the early detection of Pre-DM and T2D through gut microbiota metabolic control. They also highlight crucial targets for early intervention techniques aimed at controlling microbiome metabolism, revealing substantial predictive value and therapeutic translation potential.

### 4.4 Advantages and Shortcomings

Compared to previous meta-analyses and reviews on metabolomics and diabetes, this study is the first to employ bibliometrics to map the knowledge structure of metabolomics research related to diabetes, explicitly focusing on Pre-DM and prioritizing clinical studies. Employing various bibliometric software tools for visual analysis reveals new research themes and directions within this field, opening up additional possibilities for future investigations. However, this analysis’s potential biases could have been introduced through continuous database updates, the selection of data formats, manual screening and merging procedures, as well as the inherent algorithms of the bibliometric software used. Nevertheless, the findings of this study reflect a comprehensive bibliometric overview of research themes, hotspots, and trends in the application of metabolomics to Pre-DM and T2D.

## 5 Conclusion

Based on a bibliometric approach and perspective, this study reveals a consistent increase in the number of publications in the field of metabolomics research related to Pre-DM and T2D. Organizations, institutions, and authors from China, the United States, and Germany have made substantial contributions to the field. Metabolomics research has revealed complex metabolic disorder features associated with Pre-DM and T2D. The integration of multi-omics technologies has enabled a shift from single biomarkers to metabolic network analysis, while advances in research strategies have promoted the potential for early diagnosis and prevention of Pre-DM and T2D. Future research should prioritize elucidating the interactions between metabolism, gut microbiota, and genetic factors in diabetes, with emphasis on AI-assisted metabolic predictive models and personalized intervention strategies. Efforts are also needed to identify robust biomarkers for Pre-DM and T2D, accurately identify high-risk populations, and control diabetes. To respond effectively to address the global burden of diabetes and advance toward an era of personalized diabetic medicine.

## Acknowledgments

We thank all participants involved in this study for their time and cooperation. We also extend our gratitude to our colleagues and the research staff for their technical support and assistance during the course of this research.

